# Robust Cortical Criticality and Diverse Neural Network Dynamics Resulting from Functional Specification

**DOI:** 10.1101/2020.10.23.352849

**Authors:** Lei Gu, Ruqian Wu

## Abstract

Despite recognized layered structure and increasing evidence for criticality in the cortex, how the specification of input, output and computational layers affects the self-organized criticality has been surprisingly neglected. By constructing heterogeneous structures with a well-accepted model of leaky neurons, we found that the specification can lead to robust criticality almost insensitive to the strength of external stimuli. This naturally unifies the adaptation to strong inputs without extra synaptic plasticity mechanisms. Presence of output neurons constitutes an alternative explanation to subcriticality other than the high frequency inputs. *Degree of recurrence* is proposed as a network metric to account for the signal termination due to output neurons. Unlike fully recurrent networks where external stimuli always render subcriticality, the dynamics of networks with sufficient feed-forward connections can be driven to criticality and supercriticality. These findings indicate that functional and structural specification and their interplay with external stimuli are of crucial importance for the network dynamics. The robust criticality puts forward networks of the leaky neurons as a promising platform for realizing artificial neural networks that work in the vicinity of critical points.

## I. INTRODUCTION

With *in vitro* [1, 2] and *in vivo* [3–6] signatures across species, operating in the vicinity of a critical point is being promoted from a hypothesis [7, 8] into an established property of cortical states. Cortical activities at criticality are characterized by collective firing of neurons, known as neuronal avalanches, whose size and duration follow distributions *P*(*s*) ∝ *s*^−3/2^ and *P*(*t*) ∝ *t*^−1/2^, that is, scale-free power laws indicating a critical branching process. The criticality and scale-free dynamics have also been validated through the a renormalization group scheme for data analysis [9–11]. Functionally, it has been suggested that the capacity of information storage and processing is optimal at the critical point [12–17]. Owing to these advantages, building artificial learning models that work at the edge of phase transition is a goal of lasting efforts [18–20], and criticality is a guiding principle for devising efficient neuromorphic hardware [21–23].

As for origins of the criticality, self-organized criticality (SOC) [24, 25] is the most intensively studied theory, whereby the model may tune its parameters [26–30] according to the strength of external stimuli. Plasticity of the synaptic strength [31–33] lies at the heart of cellular mechanisms for the self-tuning. On the network level, inhibitory neurons can regulate the activity of excitatory neurons [34, 35], and their ratio is a controlling parameter for the network dynamics [36, 37]. Dynamical deployment of metabolic resources may stabilize the neuronal excitations and facilitate a critical state [38]. Structurally, adaptable network topology can foster the criticality [39]. It has been also augured that scale-free network structure [40, 41] may enhance robustness of the criticality [4, 42, 43]. The model of growing critical [44] offers an explanation from the developmental perspective. Besides criticality, it is worth noting that independent neuronal stochastic processes [45, 46] and neutral dynamics [47] can account for the observed power laws.

Despite these studies of broad scopes, how fundamental functional specification and the layered structure of the cortex influence the SOC received little attention. Here, the functional specification does not mean the opposing roles played by the excitatory and inhibitory neurons nor various cognitive functionalities of the cortical modules. Rather, it refers to different neuronal wiring patterns and specified roles of neurons therein. It is observed that certain cortical layers are main targets of input from other modules or sensory nerves, and neurons in some layers are more recurrently connected [48, 49], suggesting functional propensity for higher-order signal processing. We observed that specifying a few layers for external inputs is efficient in keeping the neural network close to the critical point, so that the dynamics is almost insensitive to the frequency of inputs. When output layers are also considered, there are both origins and destinations for the signal stream, and the distinction of feedforward and recurrent connections emerges. In this case, we found that the frequency of inputs becomes important. It may determine whether the network dynamics is subcritical, critical or supercritical.

The integrate-and-fire models [50, 51] constitute an accepted paradigm to describe the spiking dynamics of neural networks. With adaptive synapse strength, a neuron model in this line that accommodates SOC was firstly developed in Ref. [27, 28]. By incorporating membrane potential reduction due to diffusion of ions, a leaky model [30] is proposed and explains the up and down cortical states [30, 52–54]. Because of its more practical description of neuronal dynamics, we use this leaky neuron as building bricks for heterogeneous neural networks corresponding to various functional specifications. Besides the aforementioned findings, we showed that the adaptation to strong external stimuli [5] can be readily unified into this framework. Based on analytical solution of the Fokker-Planck equation, an iterative procedure of obtaining stationary state properties is developed, which is much quicker than the numerical simulation and more powerful than mean field theories, so that the heterogeneities can be addressed. Moreover, with tools in the complex network theory [55], we propose a network metric — *degree of recurrence* (DOR). Presence of output neurons reduces the DOR and leads to subcriticality. A bigger DOR can account for the more robust criticality in small-world networks [4, 42, 43] than in the Erdős-Rényi networks.

## II. ROBUST CRITICALITY

### A. The leaky neuron model

For consistency, we recap the model of Millman *et al*. [30], where each leaky neuron receives currents from and sends currents to *n_c_* other neurons, respectively. For a neuron (in terms of source neurons), the neurons being connected to (target neurons) is randomly selected. That is, the connections form an Erdős-Rényi network. Signals are transferred through vesicle release by *n_r_* release sites on each synapse. When a neuron spikes at time *t^k^*, a vesicle release is launched with a certain probability, which injects a current 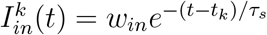 into the target neurons. Every neuron also receives independent Poisson external currents *I_e_* with amplitude *w_e_*, frequency *f_e_* and the decay time *τ_s_*. Synaptic depression is introduced in by time-dependent utility for each release site, which is set to *U* = 0 at the launching time and recovers exponentially toward 1 with characteristic time *t_R_*, i.e., *U*(*t*) = 1 − *e*^−(*t*−*t^k^*)/*τ_R_*^. When a neuron spikes, the probability of launching a vesicle release is *p_r_U*(*t^k^*), where *p_r_* < 1 is a constant and *t^k^* the spiking time.

Putting these ingredients together, the membrane potential of the ith neuron obeys the equation

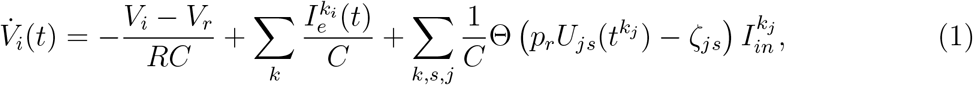

where *R* is the membrane resistance, *C* the capacitance, Θ denotes the step function, and *ζ_js_* is a random variable uniformly distributed on [0, 1]. Index *j* denotes source neurons of this neuron, and *s* indexes release sites on the synapses thereof. Upon reaching the threshold voltage *V_θ_*, the neuron spikes. After a short refractory period *τ_rp_*, the *V_i_* is reset to the resting potential *V_r_*, and during this period dynamics of the neuron is turned off. We used the parameter values in Refs. [30] and [47], with an exception *p_r_* = 0.3. Since the temporally alternative up and down states [30, 52–54] in the exact critical regime is not the major concern here, we used this value that is bigger than 0.25 to slightly tune the dynamics toward the supercritical regime.

### B. Networks with a specified input layer

To mimic the specification of input neurons and computational neurons, we simulated dynamics of a 10-layer network with 81 neurons in each layer, which is basic setting throughout this work otherwise specified. Only the neurons in the first layer receives external inputs and all the neurons are randomly connected [Fig. 1(a)]. Essentially, because of the random connections, the network does not have a layered structure. The only difference with the plain model is the specification of 1/10 of the neurons as receptors of external stimuli.

**FIG. 1.**
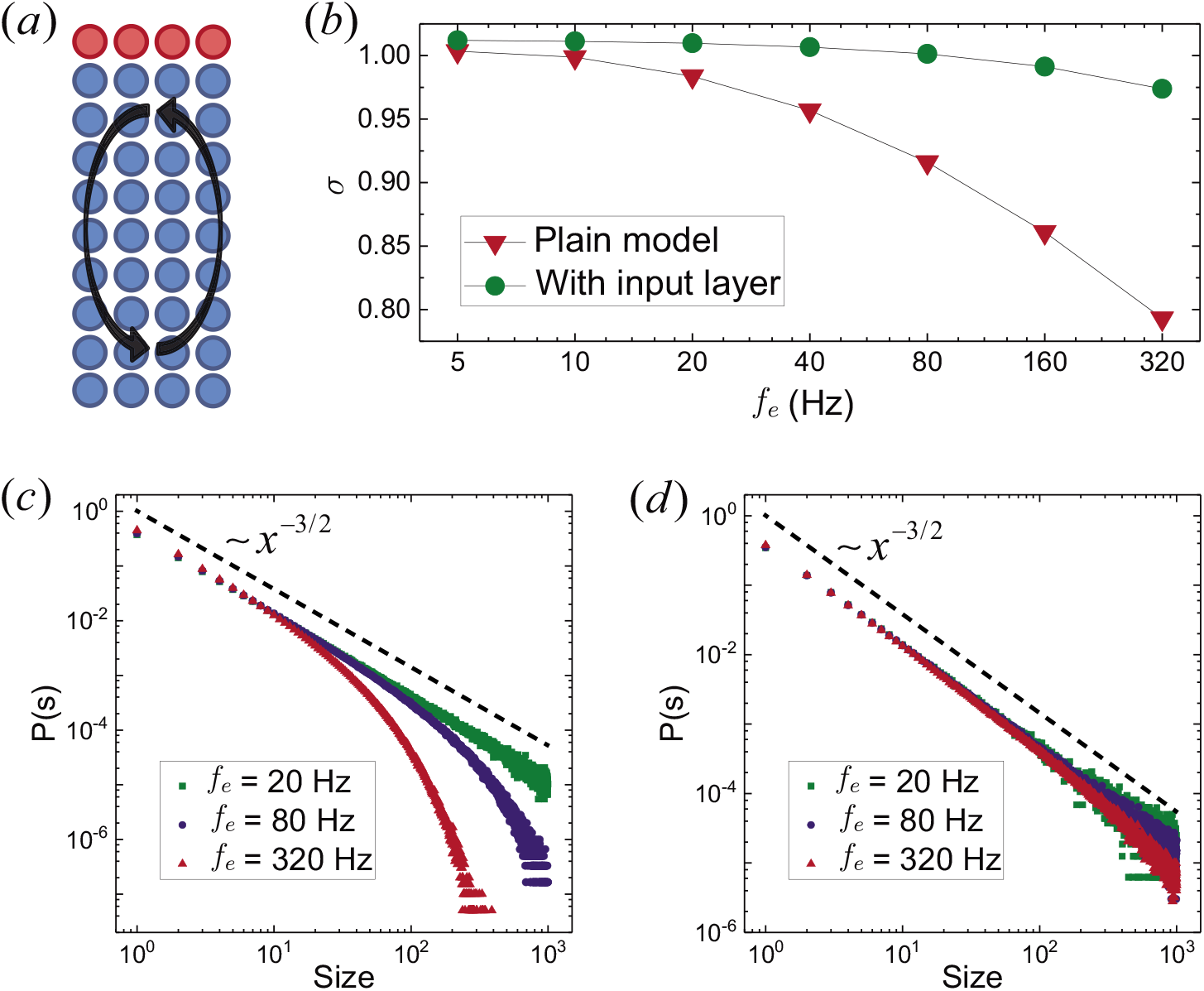
(a) Neurons in the first layer is specified as receptors of external inputs. (b) The branching rate of the network is only reduced a little by increasing frequency of inputs, while the reduction is significant for the plain model. On the contrast to distributions of the avalanche size for the plain model (c), the distributions for the case of a specified input layer (d) are close to *P*(*s*) ∝ *s*^−3/2^, demonstrating robustness of being in the vicinity of the critical point.

It is known that the unit branching rate is an indication of the critical branching process. Fig. 1(b) compares the average branching rate of the plain model and the network with a specified input layer. It shows that the specification of input layer results in a branching rate close to 1 even under high frequency inputs. This robustness of being close to the critical point can also be seen from the distribution of avalanche size. As shown in Fig. 1(d), the distribution still deviates little from *P*(*s*) ∝ *s*^−3/2^ even for inputs of frequency *f_e_* = 320 Hz. This robustness is understandable from the neuronal dynamics. When the external inputs have high frequencies, the firing rates of the recipient neurons are also high and the synaptic utility cannot sufficiently recover. This decrease in vesicle transmission efficacy reduces the ability of inducing a consequential spike in neurons they connect to. When only one layer is specified for external inputs, this suppression effect is largely relieved.

In return, the robust criticality implies a primary advantage or requirement for functionality of the cortex. As only the sensory neurons receive external inputs, and signals in the other parts of the brain are endogenous, sustainment and propagation of these input signals are prerequisites for them to reach deep regions of the cortex. A branching rate close to 1 makes such deep propagation possible in the first place, besides facilitating other functional advantages [12–17].

### C. Layered structure

To further investigate the reason for the robust criticality, we construct a layered structure as sketched in Fig. 2(a). The setting is the same as in the Fig. 1(a), except that now neuronal connections are strictly layer-to-layer. Namely, a neuron sends signals only to neurons in the next layer below it. To maintain the recurrent nature, the last (10th) layer is connected back to the first layer.

**FIG. 2.**
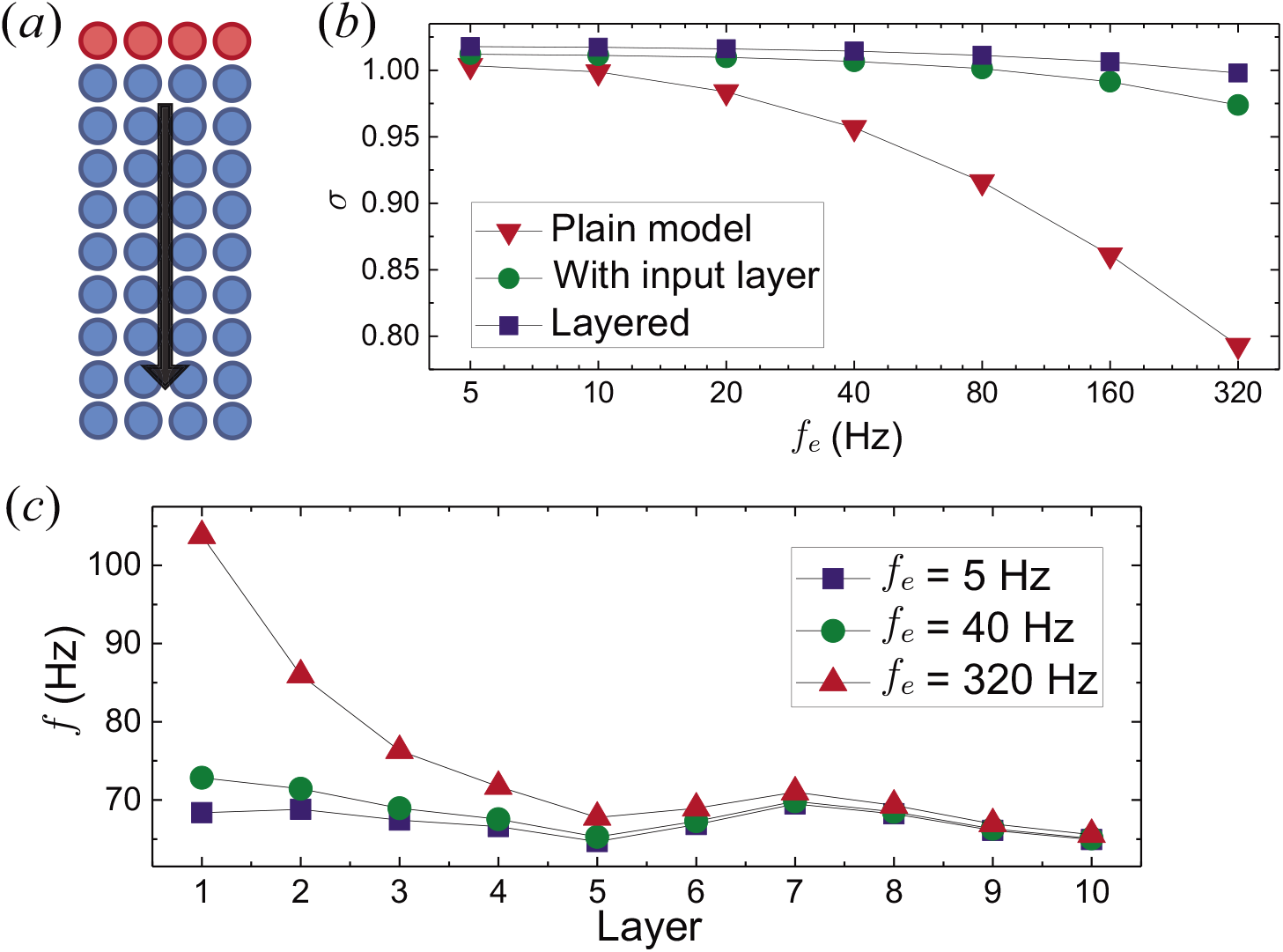
(a) In this network, the input signals propagate layer by layer. (b) Reduction of the branching rate by increasing input frequency is approximately halved compared to the setting in Fig. 1(a), indicating further enhancement of the robustness. (c) For high frequency inputs (320 Hz), when going into the inner layers, the firing rate shows exponential-like decrease and approaches its *intrinsic* values.

From the branching rates in Fig. 2(b), we see that the layered structure further facilitates the robust criticality, since the reduction caused by the increasing input frequency is approximately halved. High frequency external stimuli yield high firing rate in the input layer. However, Fig. 2(c) suggests that these stimulations only have limited effects, and the firing rate decreases following an exponential-like law. On contrary to the plain model, the firing rate for *L* > 4 in the layered network is insensitive to the inputs. This allows the identification of *f* ≈ 70 Hz as the intrinsic firing rate for neurons in this network. In other words, the reason for the robust criticality lies in the fact that the over-firing due to high frequency inputs quickly diminishes after several steps of propagation and an intrinsic firing rate is approached.

### D. Stationary state dynamics

Because of the heterogeneities, the mean field theory and stationary state results for homogeneous networks are not enough, and more advanced techniques are needed. To this end, we analytically solve the Fokker-Planck equation for each neuron to obtain relations among the neuron-wise variables, and develop an iterative procedure. Assuming that spikes of each neuron are independent Poisson processes, distribution of the membrane potential obeys the Fokker-Planck equation

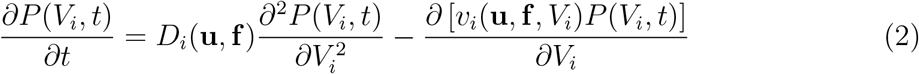

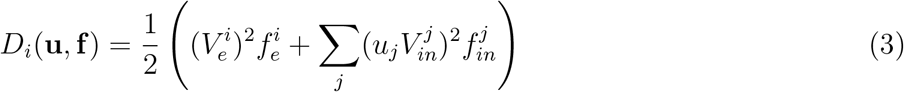

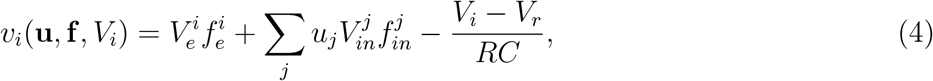

where 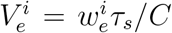 and 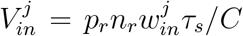. Vector **f** contains frequency of the external stimuli upon this neurons and firing rate of the neurons sending vesicle to this neuron, and elements of **u** are the mean values of the synaptic utility for these source neurons, *u_j_* = 〈*U_js_*〉.

Setting *∂P*(*V_i_, t*)/*∂t* = 0, the stationary state solution can be obtained as (Appendix A)

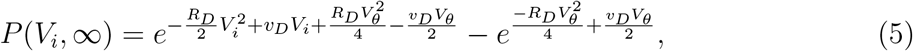

where *R_D_* = 1/*RCD_i_* and 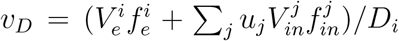. The firing rate is probability current that passes through the threshold, and thus can be derived according to 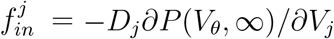. Together with *u_j_* = 1/(1 + *p_r_τ_R_f^j^*), the equations constitute a network of relations among the neuron-wise firing rates and synaptic utilities, for which the self-consistent solution can be obtained via iteration. Then, using the clever approximation (Eq. (9)–(11) therein) in Ref. [30], the branching ratio is given by

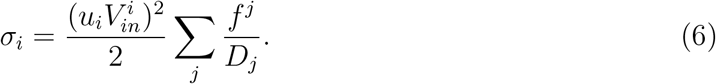

Here and on the above, Σ_*j*_ is over the source neurons.

In the layered structure, only layer-wise parameters and variables are needed. As a test of this procedure, in Fig. 3 we compare the analytical firing rate and branching rate with the numerical simulation results. In general, they agree with each other, but the firing rates in the deep layers differ a lot. The most likely reason for this discrepancy is the assumption to obtain Eq. (2)–(4) that spikes of different neurons are independent Poisson processes. This assumption is more pertinent for the layers close to the input layer, as the spikes therein are strongly affected by the external stimuli, which are independent Poisson processes. In contrast, firings in the deep layers belong to long spike trains. Induced sequentially, these firings are endogenous and causally correlated [47, 56].

**FIG. 3.**
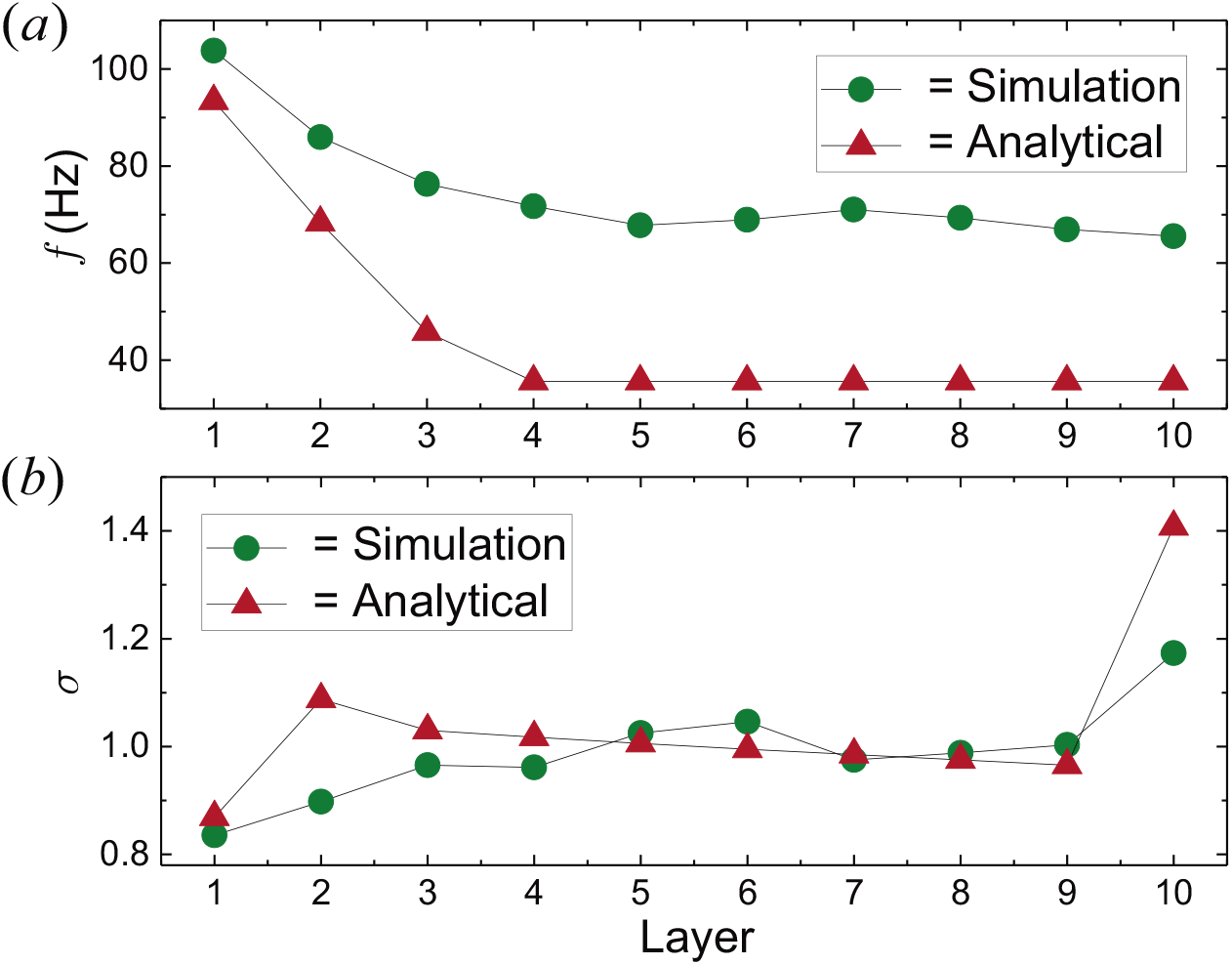
Comparison of layer-wise (a) firing rate and (b) branching rate obtained by numerical simulation and analytical solution, respectively. The disparity of the firing rate for the deep layers is likely due to the inherent correlation among the spikes, which violates the independence assumption for deriving the Fokker-Planck equation.

## III. PHENOMENAL IMPLICATIONS

### A. Adaptation to strong external inputs

In Ref. [5], visual stimuli were presented to turtles at rest. It is observed that their visual cortex adapts to the stimuli and approaches a critical working state after a transient period of burst of neuronal firings. The theoretical description there adopted a general integrate-and-fire model. The approaching to criticality relies on an updating rule such as 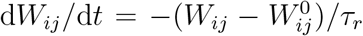, where *W_ij_* is the synaptic strength between the *i*th and *j*th neurons. 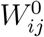 denotes prescribed strengths corresponding to the critical state and *τ_r_* is the adaptation time scale. While this model successfully describes the response of neuronal activity and synaptic strength to the stimuli, the adapted critical state is somewhat an artifact due to the prescribed parameters.

The model in Ref. [5] is homogeneous and every neuron receives external stimuli. However, we reasoned that this may not be a practical description, as only the neurons connected to the retina directly receive external stimuli. Accordingly, we construct a network architecture as shown in Fig. 4(a). The input layer mimics the retina neurons, the following three layers with feed-forward connection represent the intermediate neurons to the virtual cortex, and then layers of recurrently connected neurons model the virtual cortex. We set the frequency of the stimuli as *f_e_* = 320 Hz. Similar to the setting in Fig. 2(a), here the buffering layers (orange) ensure that the firing rate in the cortex region (blue) is sufficiently relaxed and the stationary cortical state is critical.

**FIG. 4.**
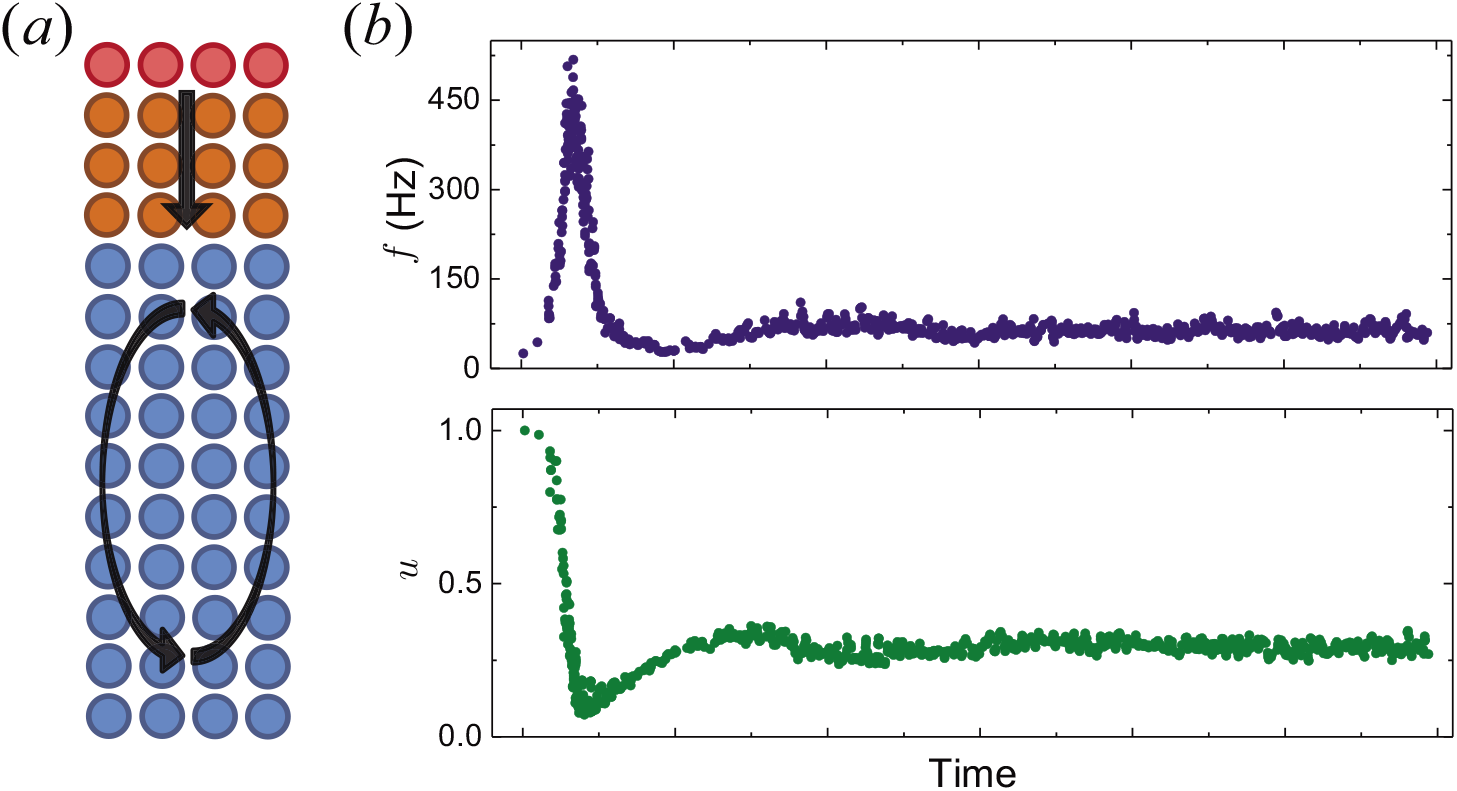
(a) Mimic of the visual pathway consists of input layer (red), transmission layers (orange), and the visual cortex neurons (blue). Evolution of the firing rate (upper) and synaptic utility (lower) of the cortical neurons is very similar to the results in Ref. [5], suggesting adaptation activity consistent with the experimental observation.

Fig. 4 (b) illustrates response of the average firing rate and synaptic utility of the cortical neurons (i.e., the blue part in Fig. 4(a)), when the frequency of inputs change from 4 Hz to 320 Hz. Considering the synaptic utility as instant strength of the synaptic connections, these curves is quite similar to the results in Ref. [5]. In other words, once we take the visual pathway as a heterogeneous structure like shown in Fig. 4(a), criticality of the virtual cortex can be readily maintained, and the adaptation activity is consistent with the experimental observation. Notably, here we only devised the wiring pattern of the neurons, and no special parameter or additional tuning mechanism is introduced in.

The key for understanding these results is the recovery dynamics of the synaptic utility, which follows *U*(*t*) = 1 – *e*^−(*t*−*t*_0_)/*t_R_*^ in the interval between two spikes. When the frequency of inputs is low, the cortical neurons are rarely excited, so *U* = 1 for most neurons. Namely, the vesicle release sites are fully powered and ready for launching. As a result, when external inputs are suddenly strengthened, the cortical neurons connected to the transmission neurons (orange) is effectively excited and their firings cause a burst of spikes in the visual cortex (blue). In the stationary state, regular spikes prevent the synaptic utility from full recovery, so the utility and firing rate are both kept at a relatively low level.

### B. Subcriticality due to output neurons

Because frequent firings of a neuron suppress the synaptic utility, high frequency of external inputs can lead to subcriticality (cf. Fig. 1). This cause of subcriticality also presents in other models [6, 27, 28, 49]. As demonstrated in the preceding, however, in a functionally heterogeneous network the external inputs only increase the firing rate of nearby neurons, and most neurons are activated in an intrinsic rhythm. This casts doubt on high frequency inputs as an explanation for the subcritical states [5, 6, 49, 57].

In contrast with input neurons, a cortex module should also send out processed signals for communication with other modules or for controlling. There much be some neurons that function as output neurons. Since the outgoing signals are not fed back, spike trains may be terminated at these neurons. At the macroscopic level, the propagation of spikes is random [47], so long spiking trains are more likely to reach the output neurons. As a result, the termination of spikes due to output neurons reduces the probability of forming large avalanches, and the dynamics appears to be subcritical.

To test this intuitive prediction, we further set the last layer of the network in Fig. 1(a) as the output layer and get the network in Fig. 5(a). Namely, we construct a network of 810 neurons that are randomly connected to (on average) 7.5 other neurons, and specify 81 neurons for inputs and outputs, respectively. The frequency of the inputs is *f_e_* = 20 Hz. To investigate how signal feedback affects the network dynamics, we tuned the number of connections from the output neurons (green) to the others (red and blue). As expected, in Fig. 5(b) the distributions of avalanche size for low degrees of feedback (*N_b_* = 0,0.25) show clear signature of subcriticality. When *N_b_* = 7.5, a fully recurrent network is recovered and the criticality is regained.

**FIG. 5.**
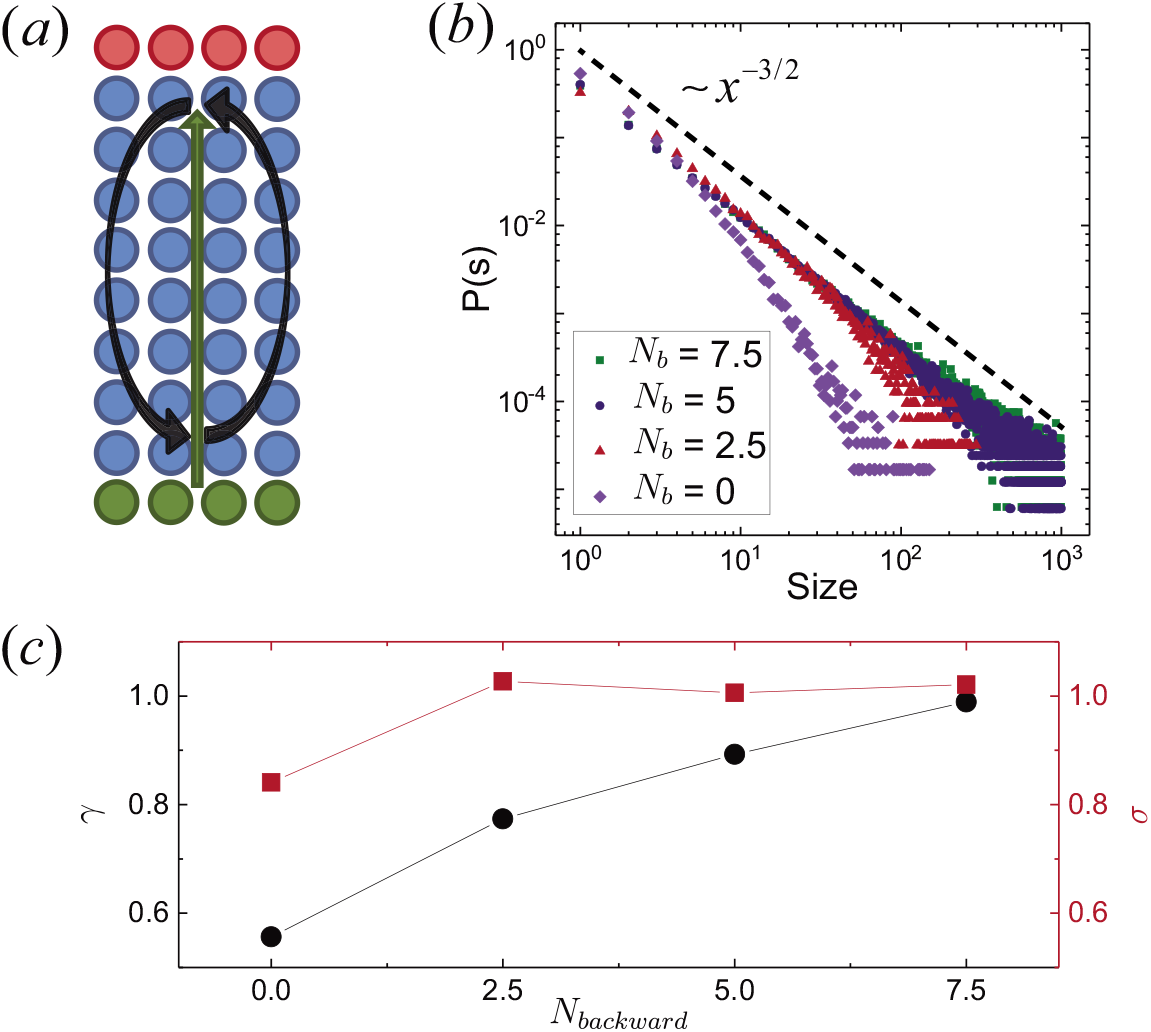
(a) A recurrent network with none, partial, or full feedback connections from the output layer. (b) Low degrees of the feedback connections leads to subcritical dynamics. (c) The degree of recurrence (*γ*) is a more pertinent metric for the network dynamics than the branching rate (*σ*); its trend is consistent with the avalanche size distributions in (b).

Because now the signal termination due to the output neurons is the major reason for subcriticality, the branching rate [*σ* in Fig. 5(c)] is not appropriate to account for it. For instance, the branching rate for *N_b_* = 2.5 is slightly bigger than 1, but the avalanche size distribution is subcritical. Based on the understanding of the cause, we propose *degree of recurrence* (DOR), which is defined as

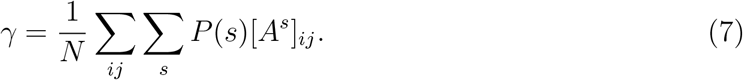

Here *A* is the adjacency matrix with row-wise normalization, so that the elements represent probability of inducing a spike in the target neurons by firing of the source neurons (see Appendix B for details). *N* is the number of neurons in the network and *P*(*s*) is the distribution of lengths of spike trains, for which we used *P*(*s*) ∝ *s*^−3/2^. Taken together, the summation is a description of how likely a spike train is sustained by the network. As shown in Fig. 5(c), the fully recurrent network has *γ* ≈ 1, and reducing the feedback connections lowers the DOR. The trend of DOR is consistent with the avalanche size distributions in Fig. 5(b).

We can also define a local version of DOR by taking a diagonal value of *A^s^, γ_o_* = Σ_*s*_ *P*(*s*)[*A^s^*]_*ii*_. It represents the probability of a spike train revisiting its origin neuron (here the *i*th neuron). In Appendix B, we show that small-world networks have bigger local DORs than the Erdős-Rényi networks. This result does not depend on the specific neuronal dynamics. Basically, it reflects the fact that spike trains in small-world networks are dispersed more slowly. In the subcritical regime, more spikes within a small region make it more likely to form collaborative excitation and helpful for sustaining large avalanches, so that the robustness to staying near the criticality [4, 42, 43] is enhanced.

### C. Diverse network dynamics

With the specification of input and output layers, a direction of the signal stream is defined, and the distinction of feed-forward and feedback connections arises. To study how these connections influence the network dynamics, we construct networks in Fig. 6(a). Besides designating the input and output layers, we specify a part of the connections of the remaining neurons (blue) for forward layer-to-layer connections (cf. Sec. II C), and the rest part for random (recurrent) connection with 7.5 connections/per neuron in total. We note that because of the random selection of target neurons, the latter part also contains forward connections, especially for the neurons near the input layer. We did not perform the detailed statistics and still count them as recurrent connections.

**FIG. 6.**
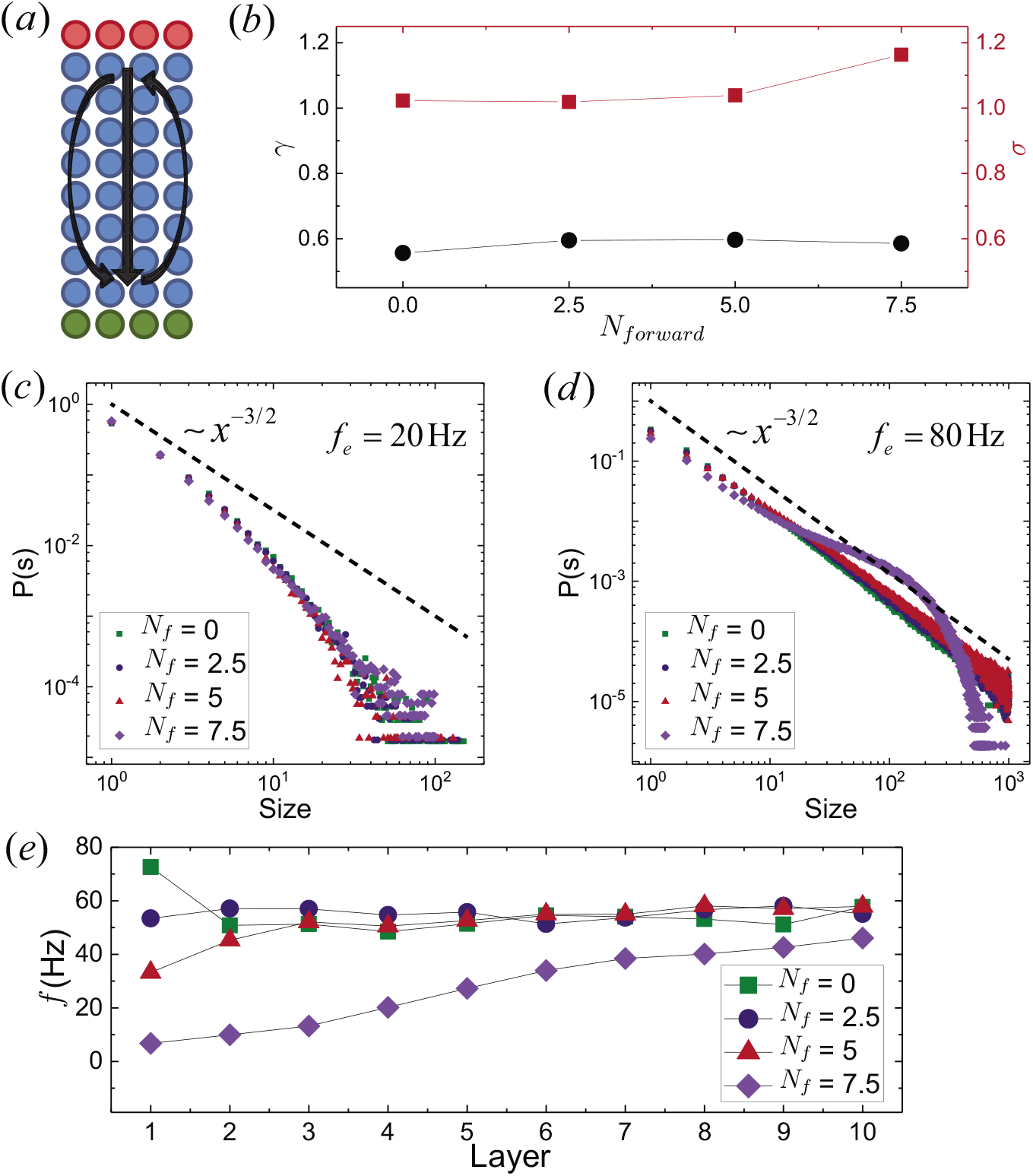
(a) Besides specification of input and output neurons, the ratio of the feed-forward and recurrent connections are varied to study its influence. (b) The branching rate for *f_e_* = 80 Hz is consistent to the avalanche size distribution in (d), where critical and supercritical dynamics is driven. But the DOR can only account for the subcriticality in (c), implying that more advanced network metrics are needed. (e) Feedback connection is a tuning mechanism for branching; when they are insufficient, the dynamics may be driven to the superciricality by high frequency inputs (here *f_e_* = 80 Hz); this is indicated by the monotonically increasing firing rate, implying branching rate *σ* > 1 in each layer.

Fig. 6(c) suggests that low frequency inputs can only lead to subcritical network dynamics, and ratio of the two types of connections matters little to the avalanche size distribution. As shown in Fig. 6(d), however, high frequency inputs can drive the dynamics close to the criticality. When the ratio of feed-forward connections is sufficiently high, supercritical dynamics can be driven, as suggested by the arc for *N_f_* = 7.5. The critical and supercritical dynamics is further indicated with the branching parameter (*σ*) in Fig. 6(b). Fig. 6(b) also gives the DOR, which is a network property independent of *f_e_*. While these *γ* < 1 can characterize the subcriticality for low frequency inputs, they cannot account for the cases of high frequency inputs. An input-dependent metric is required, which we leave for further study.

These diverse dynamics imply interplay of external stimuli and the two types of connections. In the first place, sufficient stimuli are required to sustain large avalanches, or otherwise the dynamics is subcritical. The firing rate curves (here *f_e_* = 80 Hz) in Fig. 6(e) suggest that the feedback connections in itself is a tuning mechanism. Namely, they suppress the branching and keep the branching rate close to 1. Otherwise, the branching rate can be considerably bigger and the dynamics is driven to the supercritical regime. This is clear in the firing rate curve for *N_f_* = 7.5. Without feedback connections, the firing rate increases monotonically from the input layer to the out layer, indicating an branching rate *σ* > 1 in every layer. Besides input strength [49], these results suggest connectivity pattern as a cause of various network dynamics.

## IV. DISCUSSIONS AND CONCLUSION

Being an intricate information processing system, the cortex together with other parts of the brain receives, transmits, functionally processes, and sends out signals. The homogeneous networks for studying the cortical criticality are an over-simplified description. Smallworld and scale-free networks are endowed with certain wiring patterns of the neurons, but still not enough to account for the rich functional and structural specifications. Our results demonstrate that these specifications and their interplay with external stimuli are important in determining the dynamics of the neural network. Accordingly, existing understandings of the network dynamics should be reconsidered, especially their network-structural origins. The network structures here are prototypical, and more delicate neuronal wiring may be need to describe specific parts of the brain. Other problems such as effects of inhibitory neurons and other self-tuning mechanisms [31] are interesting and worth investigation. Containing only the structural information, the *degree of recurrence* is in its primitive form, and substantial extension may be required to deal with the various influential factors.

Despite lasting attempts to construct neural network models that can carry out cognitive tasks and meanwhile work in the critical state, limited successes are achieve. An outstanding issue is the unmaintainable criticality due to input signals—the network at rest can be in a critical state, but the criticality is compromised by signal inputs [18, 20]. Several factors in the past endeavors could hinder the success. One is the choice of neuronal models. For instance, when models with fixed synaptic strength are used [18], self-tuning mechanisms play no role. Moreover, a recent study suggests that some of the used model may not describe the branching process but random walk [58]. Another aspect is how to use the network models. There is a paradigm whereby the network is used as an encoding apparatus of the input signals [18, 59], and the functional specification is not well leveraged. Regarding these possible hindrances, the robust criticality with respect to external stimuli and the input driven criticality provide valuable insights for building artificial intelligence networks that have critical working states.

In conclusion, by constructing various heterogeneous network structures with the leaky neuron, we achieved output-induced subcrticality, robust criticality, input driven criticality and supercriticality, and criticality ensuing transient adaptation to external stimuli. An analytical procedure and network metrics are proposed to account for the heterogeneities. The diverse dynamics put forward many topics about how neuronal dynamics, connectivity patterns and external stimuli affect dynamics of the neural network. It is also intriguing to develop networks of the leaky neurons into models that are capable of learning. This may provide a generic route to artificial intelligence networks that work in the vicinity of criticality. Endowed with neuronal dynamics accommodating criticality, such models can also shed practical light on how learning shapes properties of the cortex.

## ACKNOWLEDGEMENTS

We are grateful to Jorge Hidalgo for valuable suggestions and discussions.

## Appendix A: Solution of the voltage distribution

Before the derivation, a possible puzzle of the Fokker-Planck equation is worth noting. In Eq. (2)–(4), the *V*_i_ is a variable specifically for the ith neuron, and not position-dependent. How network structure affects the neuronal dynamics are reflected by the fact that *P*(*V_i_*) is a neuron-wise function, that is, it varies neuron by neuron. We should also note that the rescaling procedure of the incoming currents in Ref. [54] was applied to the analytical results, with reduces the disparity with simulation results.

In the stationary state, the Fokker-Planck equation Eq.(2)–(4) is in the general form,

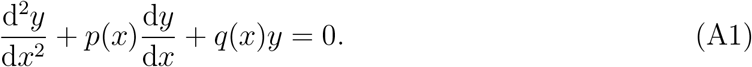

Assuming a variable separation *y* = *U*(*x*)*v*(*x*), the equation amounts to

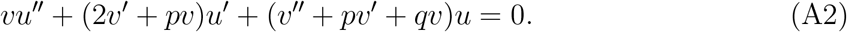

The second term can be eliminated by *v* taking the form 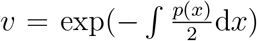. Then, the equation reads

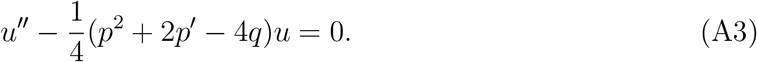

With variable notation *x* = *V_i_* – *V_r_*, in our case *p*(*x*) = *R_D_x* – *v_D_* and *q*(*x*) = *R_D_*, where *R_D_* = 1/*RCD_i_* and 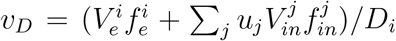. Regarding the quadratic factor of *u*, we tried solution *u* = (*ax* + *b*)exp(*cx*^2^ + *dx*). By comparing the coefficients, we obtain two solutions of *u*, which are 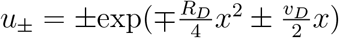.

The coefficients of the two special solutions (i.e., *vu*±) are determined by the boundary condition and normalization. As the voltage is set to *V_r_* once it reaches the threshold *V_θ_*, *P*(*V_θ_*) is virtually zero, leading to the boundary condition *P*(*V_θ_*) = 0. From it, Eq. (5) can be obtained. Because there is no dynamics that further lowers the voltage when *V_i_* = *V_r_, V_r_* is the lower limit of the voltage, and the normalization constant is determined by

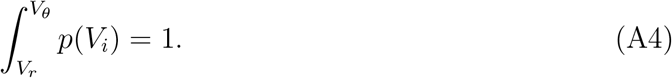

With variable change *V_i_* = *x* = *V_i_* – *V_r_*, the integration interval is [0, 20] mV for Eq. (5).

## Appendix B: Degree of recurrence

In the network in this article, the connection is directed. The adjacency matrix consists of 1s and 0s. *A_ij_* = 1 indicates that there is a connection from the *i*th neuron to the *j*th neuron, or otherwise the element is zero. With the row-wise normalization (divided by *n_c_*), the elements represent the probability of inducing a spike in the target neurons by firing of the neuron. Without random fluctuation of the number of connections, which is impossible for a fraction such as *n_c_* = 7.5, sum of every row is exactly normalized to 1. Then, for a fully recurrent network

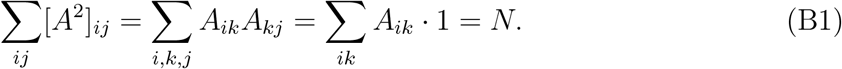

Similarly, Σ_*ij*_ [*A^s^*]_*ij*_ = *N* for *s* > 2. So, if *P*(*s*) in the definition Eq. (7) is a normalized probability distribution, we have *γ* =1. Randomness in the number of connections for each neuron may cause small deviation from the unity. For neurons in an output layer, ∑_*j*_ *A_ij_* < 1 and accordingly *γ* < 1 for networks with output neurons.

Taking the local value of the degree of recurrence, we have for the *i*th neuron

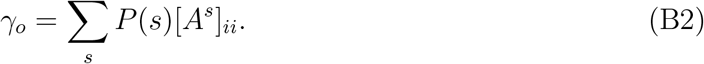

It is the probability of a spike train revisiting the *i*th neuron that is weighed by the size distribution of avalanches. In a network where a neuron connects with the other neurons with equal probability, the probability of reaching a neuron does not depends on the propagation step and follows uniform distribution. Namely, [*A^s^*]_*ii*_ is independent of *s* and 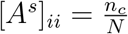.

For small-world networks, we consider a 2D network where the connection probability follows the Gaussian distribution. That is,

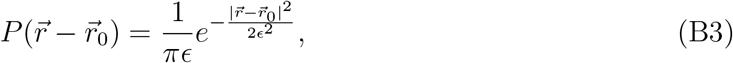

where 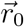 and 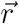 are positions of a source and its target neurons, and *ϵ* denotes the standard error. In every propagation step, the probability of generating a spike at the *j*th neuron is the probability of firing of the *i*th neuron times the connection probability from neuron *i* to *j*. Namely, the distribution of spikes in a spike train satisfies a recursive relation

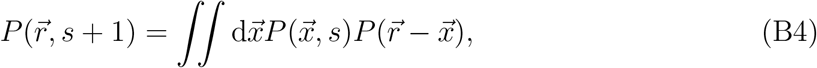

where 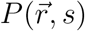 denotes step-dependent distribution of spikes in a train and should be distinguished with the connection probability 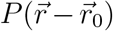. We somewhat abuse the index *s* and this may blur the distinction among avalanche size, spike train length and propagation steps, but for a branching rate close to 1, they take similar values. By Fourier transformation and the convolution theorem, we can deduce that

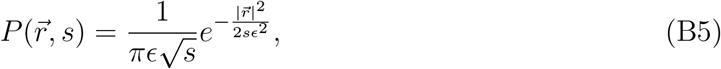

that is, a Gaussian distribution with the squared error linearly expanded in every propagation step.

In the derivation, we take the network as continuous media of spikes. Going back to discrete representation, the probability of the spike train revisiting the origin at step *s* is *P*(*s, r*) integrated over the unit area around the origin. Namely, we have

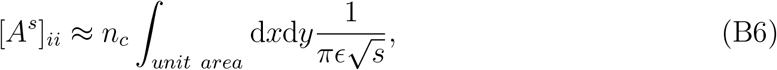

where *n_c_* arises since 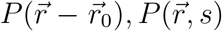 are normalized probability, but there are *n_c_* connections for every neuron. The large step limit of 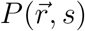 is the uniform distribution, so [*A^s^*]_*ii*_ = *n_c_*/*N* for *s* → ∞. For every finite propagation step, we have [*A^s^*]_*ii*_(*s*) > *n_c_*/*N*. Therefore, the small-world networks have local DORs bigger than the Erdőos-Réenyi networks. By virtue of the Gaussian form, the derivation can be readily generalized to the cases of other dimensionalities. Basically, the bigger magnitude of the small-world network is because the connection probability near the origin is denser than the uniform distribution. Distributions other than the Gaussian may not lead to an analytical expression for 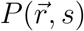, but the above argument is supposed to have general viability for other distribution of the small-world feature.

